# Evolutionary divergence in sugar valuation shifts *Drosophila suzukii* oviposition choice towards ripe fruit

**DOI:** 10.1101/2022.06.22.497171

**Authors:** Matthieu Cavey, Bernard Charroux, Solène Travaillard, Gérard Manière, Martine Berthelot-Grosjean, Sabine Quitard, Caroline Minervino, Brice Detailleur, Yaël Grosjean, Benjamin Prud’homme

**Affiliations:** Aix-Marseille Université, CNRS, IBDM, Institut de Biologie du Développement de Marseille, Campus de Luminy Case 907, 13288 Marseille Cedex 9, France; Centre des Sciences du Goût et de l’Alimentation, CNRS, INRA, Institut Agro, Université de Bourgogne Franche-Comté, 21000 Dijon, France

## Abstract

Behavior evolution can promote the emergence of agricultural pests via ecological niche changes. The underlying neuronal mechanisms are poorly understood. We investigate the chemosensory changes underlying the evolutionary shift of oviposition substrate of the pest *Drosophila suzukii* from rotten to ripe fruits. Using a model substrate for fermented fruits and genetic manipulations, we show that an increase in valuation of fruit sugars during oviposition decisions drives *D. suzukii* to oviposit on ripe fruits as opposed to the model *D. melanogaster* which prefers rotten fruits. Inter-species comparative *in vivo* calcium imaging of sugar Gustatory Receptor Neurons suggests that increased sugar valuation in *D. suzukii* is related to neuronal sensitivity changes at multiple levels of the oviposition circuitry. Our results show that the tuning of sugar valuation has contributed to the evolution of oviposition preference on ripe fruit of *D. suzukii*.

## Introduction

The emergence of agricultural pest species can occur via morphological and/or behavioral adaptations enabling access to new ecological niches. Understanding the evolution of a novel pest species therefore entails to identify the presumptive genetic, molecular and cellular modifications underlying this type of adaptation. Insect sensory-driven behaviors - in particular chemosensory behaviors – have proven highly valuable to study how a given species chooses and adapts to a particular niche. Chemosensory information guides feeding, reproduction and social behaviors. The molecular signature of natural substrates (e.g. fruits or mates) is decoded by neuronal circuits receiving information from olfactory and gustatory sensory neurons expressing large families of chemoreceptors each tuned to a range of specific molecules (reviewed for e.g. in ref. ^1^).

Evolution of behavior has often been studied in the context of specialization on a given host for feeding, ovipositing or specific mates for reproduction. In most examples, host specialization is promoted by tuning the behavior to stimuli that are specifically present in the host and frequently involves changes occurring at the level of the sensory neurons detecting these molecules ^2–12^. Changes in the Central Nervous System (CNS) affecting specific genes or wiring patterns have also been identified but deciphering how they contribute to behavioral shifts is more challenging ^2,13–15^. Evolutionary changes in host preference of generalist species which live and reproduce on a wide range of hosts has been less studied and whether it relies on similar strategies and molecular mechanisms remains to be determined. *Drosophila suzukii* has evolved a novel preference for laying its eggs in ripe fruits as opposed to most other *Drosophila* species such as *D. melanogaster* which prefer laying in fermented and rotten fruits. This novel behavior causes substantial damage to the fruit industry as *D. suzukii* is spreading across the world ^16,17^. *D. suzukii*’s adaptation has involved both morphological evolution of its ovipositor ^18,19^ and behavioral evolution of mechanosensory and chemosensory responses to egg-laying substrates ^20^. Chemosensory cues are particularly informative on the nature of oviposition substrates. To identify the neuronal bases of inter-species divergent chemosensory responses, we first need to identify the critical molecules present in ripe and fermented substrates to which *D. suzukii* responds differently than other species. A recent study suggests that a reduction of *D. suzukii*’s responses to bitter compounds might have relieved an oviposition inhibition on ripe substrates ^21^. Conversely, an increased response to fermentation by-products might have contributed to driving *D. suzukii* away from rotten substrates ^22^.

Here, we focus on the attractive cues that promote *D. suzukii* oviposition preference for ripe substrates and we demonstrate that fruit sugars have played a pivotal role in the behavioral divergence of this species.

## Results

### Divergent oviposition preferences can be recapitulated with a fermented strawberry substrate

The fruit maturation stages preferred by *D. suzukii* and *D. melanogaster* are loosely defined as a fruit matures gradually from unripe, to ripe, to overripe, to fermented and rotten. The lack of precise and objective definitions of, and therefore distinctions between, ripe and fermented stages limit our capacity to identify which cues are differently perceived or interpreted by different species. To produce a reliable source of fermented substrate we developed a controlled fermentation protocol of an industrial frozen organic strawberry purée -strawberry being a common target of *D. suzukii* in the wild-considered as at the “ripe” stage, inoculated with the yeast *Saccharomyces cerevisiae* and the acetic acid bacteria *Acetobacter pomorum*. In this system, the two microorganisms degrade sugar to produce fermentation products including alcohol (yeast) and acetic acid (bacteria).

We optimized inoculation conditions and fermentation duration empirically using oviposition choice behavior of wild type *D. melanogaster* and *D. suzukii* as a readout. The two species displayed opposite preferences, with *D. melanogaster* choosing the fermented substrate while *D. suzukii* choosing the ripe substrate (**Fig. 1a**). Fermentation during 3 days almost completely depletes glucose and fructose and produces acetic acid and ethanol (**Fig. 1b**). Strikingly, *D. suzukii* is one of the very few species preferring the ripe substrate among a range of closely and more distantly-related species (**Fig 1c**). Thus, our controlled fermented substrate recapitulates the novel and divergent behavior of *D. suzukii* reported previously with whole fruits ^20^. Control experiments show that the species difference in preference is not due to different responses to the sugar-rich diet required for *D. suzukii* rearing and which could have desensitized *D. melanogaster* sugar-responding GRNs ^23^ (**Supplementary Fig. 1a**). Egg-laying stimulation (no-choice) assays with only ripe or fermented substrates demonstrate that both substrates are similarly acceptable to the two species (**Supplementary Fig. 1b**). Thus, it is the relative value of these substrates to guide oviposition decision that has diverged between *D. melanogaster* and *D. suzukii*.

**Fig. 1.**
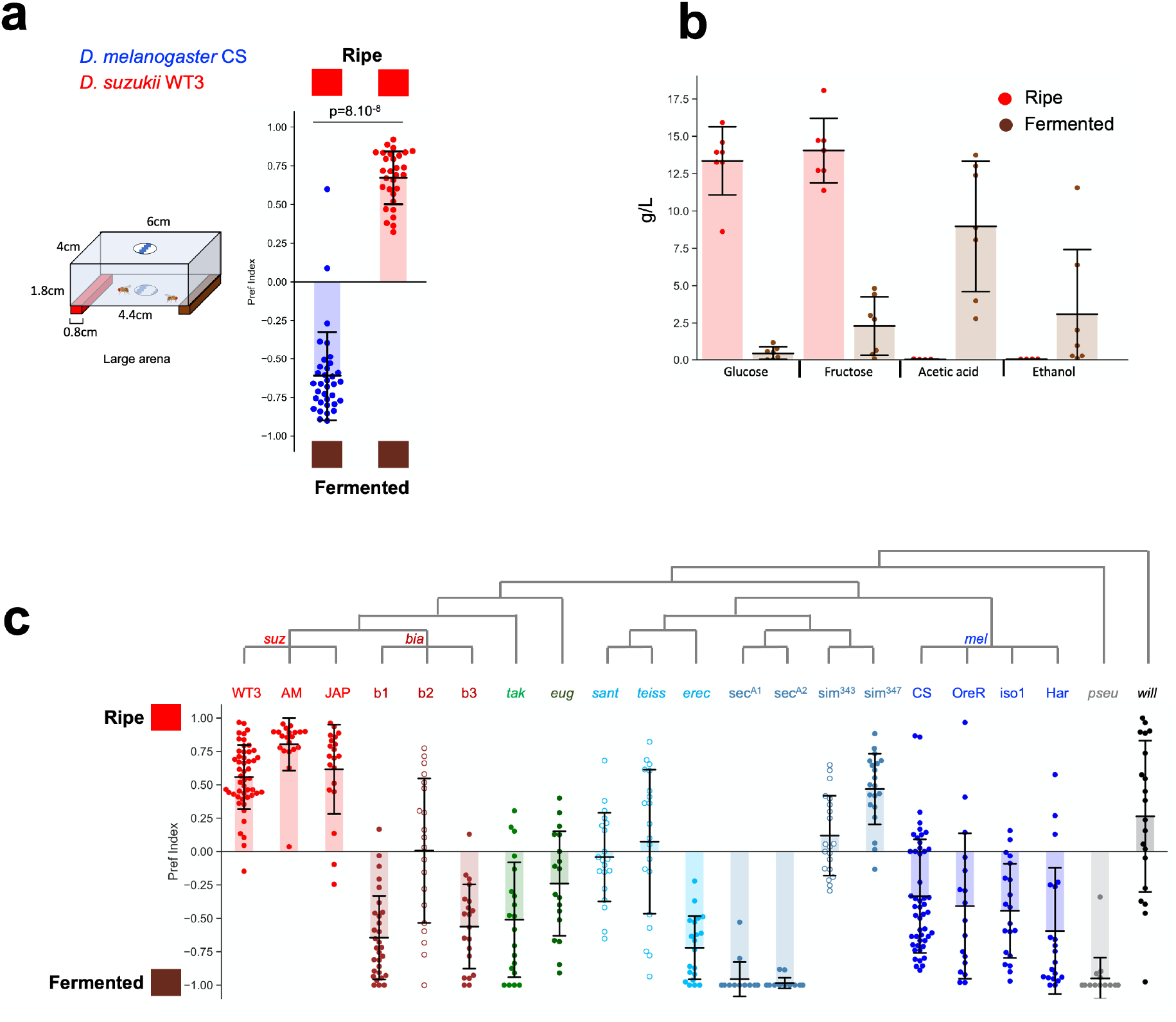
Controlled fermentation recapitulates species oviposition substrate preferences on natural substrates. **a** Oviposition substrate preference in a two-choice assay opposing ripe vs fermented strawberry purée (large assay chamber shown on left) for wild type *D. melanogaster* (blue) and wild type *D. suzukii* (red) females. The red and brown boxes on top/bottom of the graph represent ripe and fermented substrates, respectively, here and in subsequent figures. Filled circles on the graph indicate a significant preference for one of the two substrates (the distributions of preference indexes are different from zero, Mann Whitney paired test). Open circles in subsequent figures indicate no significant preference for either substrate. Shaded bars indicate the mean and error bars the standard deviation. Species preferences are significantly different from one another (Mann Whitney U test, p-value on the graph) (n=35, 30 replicas). **b** Glucose, fructose, acetic acid and ethanol levels measured in g/L (mean + standard deviation) in the ripe and fermented strawberry purées before dilution in the oviposition substrates. Fermentation effectively depletes sugars and produces acetic acid and ethanol. **c** Ripe vs fermented substrate oviposition assay for multiple *Drosophila* species. Several wild type strains were used for some species (see Methods for species names). Most species show a clear preference for the fermented substrate, apart from a few exceptions including *D. suzukii* (n=50, 20, 20, 30, 20, 20, 20, 18, 20, 20, 20, 15, 15, 20, 20, 49, 17, 20, 20, 18, 20).

In sum, controlled fermentation of strawberries produces a substrate highly attractive and stimulating to *D. melanogaster* and much less attractive to *D. suzukii* compared to unfermented strawberries. These differences recapitulate the species preferences reported for whole fruits in the wild.

### Greater valuation of fruit sugars by *D. suzukii* than *D. melanogaster* in oviposition decisions

Having established a behavioral paradigm for ripe vs fermented fruits, we went on to test whether the sugar content of these substrates contributes to species divergent oviposition preferences. Indeed, since *D. suzukii* targets many different ripe fruits in the wild, females may be stimulated to lay eggs by a cue(s) that is common to all ripe fruits. Fruit sugar (glucose + fructose) is present in all fruits, and could be a good indicator of fruit maturity since its concentration rises during maturation to peak at the ripe stage (typically ∼5-6% in strawberries ^24,25^) and then decrease upon fermentation.

To test this idea we first examined how *D. melanogaster* and *D. suzukii* value sugar as an oviposition stimulant. Previous studies have shown that the value of sugar can range from highly positive to highly negative for *D. melanogaster* depending on experimental context, in particular the arena dimension and spacing of oviposition substrates ^26,27^. We thus assessed the oviposition value of sugar-alone in two experimental setups with different dimensions. For both species, the sugar substrate (glucose + fructose at the concentration found in the ripe strawberry substrate; i.e. final concentration 1.6%) was clearly preferred against a plain agar option in both experimental setups (**Fig. 2a**), indicating that sugar-alone has a positive value for both species in our experimental conditions.

**Fig. 2.**
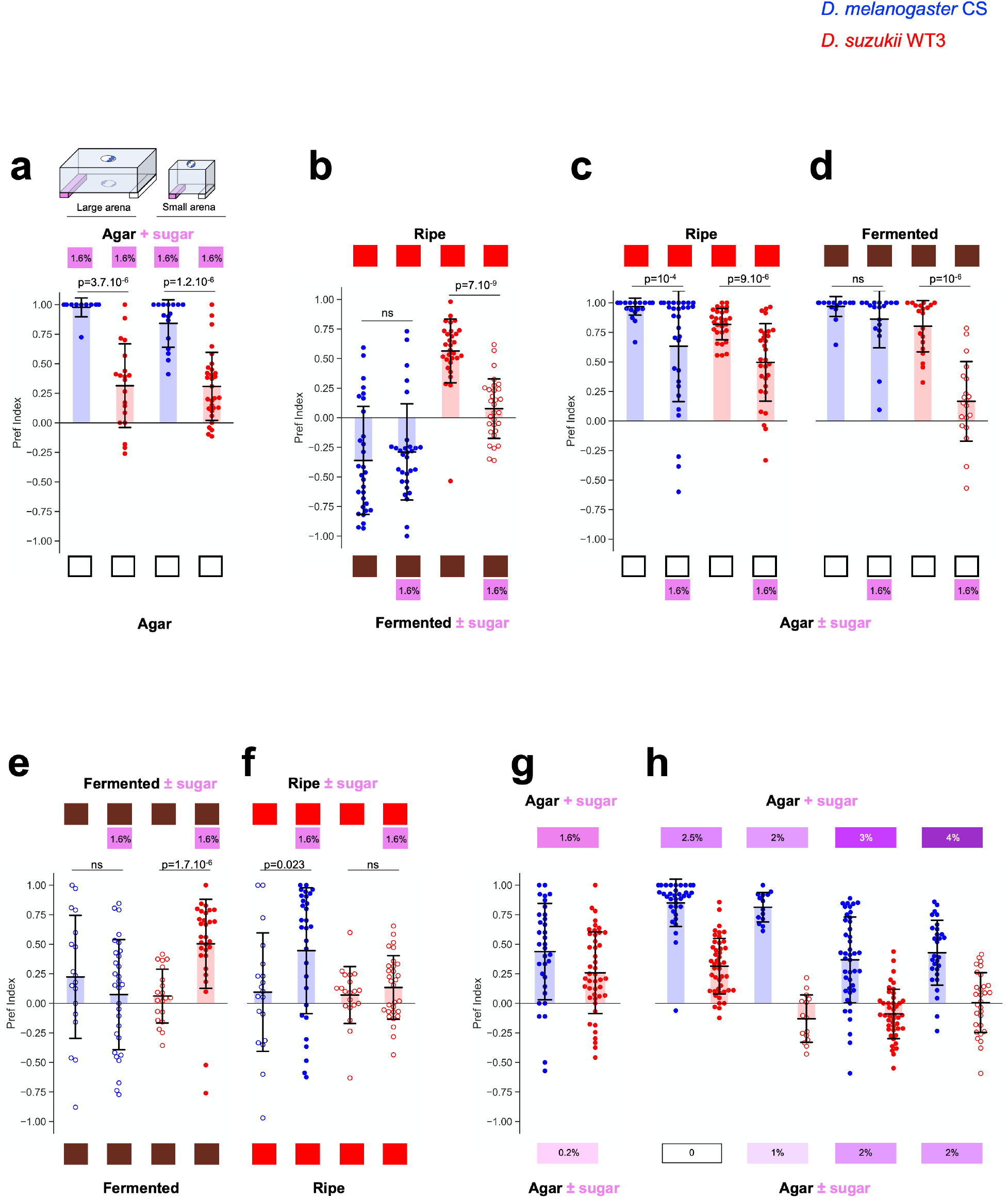
Sugar is valued differently by *D. melanogaster* and *D. suzukii* during oviposition decisions. **a** Two-choice oviposition assays opposing sugar (at the concentration of a ripe substrate, i.e.: 1.6% glucose + fructose, pink boxes on top) to plain agar (empty boxes at bottom) in the large and small chambers. Sugar is preferred by both species over plain agar in both experimental setups, but the preference is more pronounced for *D. melanogaster* (n=12, 20, 16, 30). **b** Ripe vs fermented choice assay with or without sugar added (pink box at bottom) to the fermented substrate. Adding sugar to the fermented substrate abolishes *D. suzukii*’s preference for the ripe substrate (n=30, 30, 30, 30). **c, d** Two-choice assays opposing the ripe (**c**) or fermented (**d**) substrates to plain agar or agar+sugar at the concentration of the fruit substrates. Sugar reduces the preference for ripe in both species. In contrast, sugar has almost no weight against the fermented substrate for *D. melanogaster* whereas it completely abolishes *D. suzukii*’s oviposition attraction to this substrate (n=28, 30, 29, 30, 20, 20, 20, 20). **e, f** Two-choice assays opposing the fermented (**e**) or ripe (**f**) substrates to identical substrates with or without sugar added. Sugar is valued positively by *D. suzukii* but not *D. melanogaster* in the context of a fermented substrate. Conversely, sugar is valued positively by *D. melanogaster* but not *D. suzukii* in the context of a ripe substrate (n=18, 30, 20, 30, 19, 30, 20, 30). **g, h** Two-choices assays opposing different concentrations of sugar-only. **g** *D. suzukii* is able to discriminate 0.2% vs 1.6% glucose+fructose, i.e.: concentrations matching those of the fermented and ripe substrates, respectively (n=33, 45). **h** Assays opposing different concentrations of glucose. *D. suzukii* does not discriminate concentrations ≥ 1% glucose, whereas *D. melanogaster* always chooses the higher concentration (n=35, 45, 14, 15, 43, 44, 29, 30).

We then manipulated the sugar concentration of complex fruit substrates. Adding sugar (1.6% glucose + fructose) to the fermented substrate at a concentration matching that of the ripe substrate did not change the preference of *D. melanogaster* for the fermented substrate (**Fig. 2b**). In contrast, raising the sugar concentration of the fermented substrate completely abolished the preference of *D. suzukii* for the ripe substrate (**Fig. 2b**). Thus, ripe and fermented substrates are equally attractive to *D. suzukii* as long as their sugar concentrations are equal, demonstrating that sugar is a major determinant of substrate preference for *D. suzukii*. In comparison, equalizing sugar concentration across substrates is not sufficient to give equal value to these substrates for *D. melanogaster*.

We further assessed the relative value of sugar by opposing it to either ripe or fermented substrates. Both substrates are highly preferred by each species when opposed to plain agar. However, when 1.6% sugar is added to the plain agar, the preference for ripe is significantly decreased in each species to a similar extent (**Fig. 2c**). Thus, in this context, sugar-alone appears to have a similar value for *D. suzukii* and *D. melanogaster* when opposed to a sugar-containing fruit substrate. In contrast, when opposed to sugar, the fermented substrate is still highly preferred by *D. melanogaster* but not anymore by *D. suzukii*. In fact, sugar-alone appears to be equally attractive to *D. suzukii* as the fermented substrate (preference index is not significantly different from 0; **Fig. 2d**). Thus, in this context, the relative value of sugar is far greater for *D. suzukii* than *D. melanogaster*.

Additional experiments confirmed that sugar is generally valued more highly by *D. suzukii* but this is highly context-dependent. For instance, sugar had a higher relative value for *D. suzukii* compared to *D. melanogaster* when a fermented strawberry substrate was present on both sides (**Fig. 2e**) but the two species showed the opposite trend when a ripe substrate was present on both sides (**Fig. 2f**). This last experiment suggested that as long as sugar is present on both sides - at least at a concentration ≥ 1.6% - the absolute concentration is not important to *D. suzukii*. In contrast, *D. melanogaster* appears to give more importance to the higher sugar concentration around this range of concentrations and in the absence of fermentation products.

These observations prompted us to ask whether *D. suzukii* is able in fact to discriminate the sugar concentrations present in our ripe vs fermented substrates. We thus simplified the stimuli to only sugar and opposed 0.2% glucose+fructose (∼fermented substrate) to 1.6% (∼ripe substrate). Both species chose the sweeter substrate (**Fig. 2g**), therefore, *D. suzukii* can discriminate concentrations across this range and could thus in principle use this information when making a choice between ripe and fermented substrates. Besides, these results further confirm that although the sugar content of a ripe substrate has a positive value to *D. melanogaster* when presented alone, it is outweighed by fermentation products in the context of a ripe vs fermented choice assay.

Since sugar appears to be a more potent cue for oviposition in *D. suzukii* than *D. melanogaster* in most experimental contexts, we decided to further characterize the ability of both species to discriminate small sugar concentration differences using additional competition assays with sugar-only. *D. melanogaster* consistently chose the high-sugar option, even when the low concentration substrate contained a relatively high dose of sugar (2%). In contrast, *D. suzukii* was a much poorer discriminator and did not show any preference as long as the low-sugar substrate contained ≥ 1% sugar (**Fig. 2h**). Taken together, our results are consistent with the idea that *D. suzukii* gives high value to sugar at most concentrations and in presence of complex fruit substrates, and that sugar has a low value only when present at a concentration nearing that of a fermented substrate. In contrast, *D. melanogaster* gives low value to sugar in most contexts, except when sugar is presented alone, or in the absence of fermentation products, or when presented at a relatively high concentration. Overall, this suggests that a change in sugar valuation processes could underlie *D. suzukii*’s divergent preference for ripe vs fermented substrates.

### D. suzukii is more responsive to sugar in oviposition stimulation assays than D. melanogaster

As an independent method to assess the value of sugar in oviposition decisions, we quantified the potency of individual sugars - glucose, fructose and sucrose - to stimulate oviposition in no-choice assays. While *D. suzukii* lays more eggs than *D. melanogaster* at all concentrations tested, both species respond to sugar in a dose-dependent manner, reaching a maximum egg-laying rate at intermediate sugar concentrations (**Fig. 3**). Strikingly, *D. suzukii* seems to begin responding and to reach its maximum egg-laying rate at lower sugar concentrations than *D. melanogaster*. We quantified this difference more precisely by estimating the Effective Dose 50 (ED50; sugar concentration inducing half-maximal egg-laying rate) as a proxy for the oviposition stimulation power of each sugar in the two species. The ED50 was consistently lower for *D. suzukii* compared to *D. melanogaster* (9.7-fold lower for glucose, 5.2-fold for fructose and 4.4-fold for sucrose). Thus, sugar is a more potent inducer of egg-laying in *D. suzukii* compared to *D. melanogaster*, consistent with the hypothesis that a greater valuation of sugar by *D. suzukii* biases its preference towards ripe fruits.

**Fig. 3.**
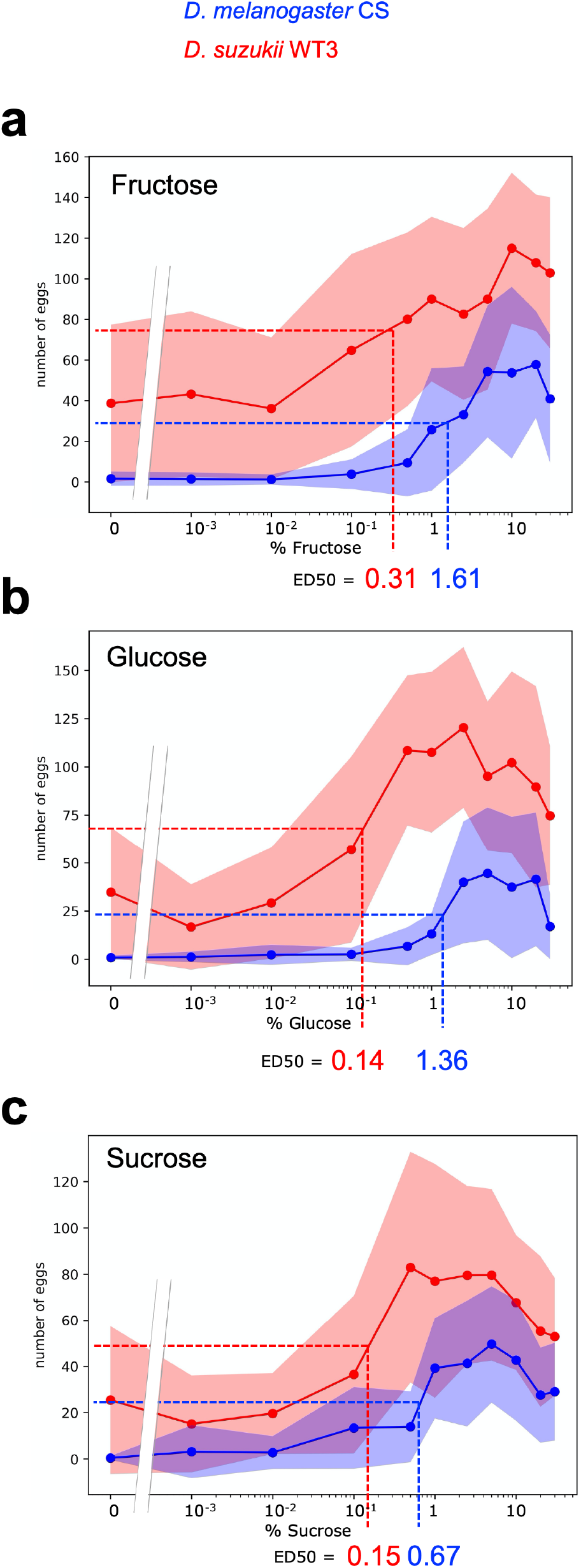
*D. suzukii* responds to lower sugar concentrations than *D. melanogaster* in oviposition stimulation assays. Oviposition stimulation assays (no-choice) on increasing concentrations of sugar-only. Egg-laying dose-response curves for fructose (**a**), glucose (**b**) and sucrose (**c**). Data is shown as the mean (dots+lines) +/-standard deviation (shaded areas). The sugar concentration (x-axis) is in log scale. The estimated ED50s are indicated at the bottom and with the dashed lines. The ED50 is consistently lower for *D. suzukii* compared to *D. melanogaster* (n=30 for each condition).

### Sugar perception is required for oviposition decisions in *D. suzukii*

To formally test this hypothesis, we developed genetic tools to manipulate sugar perception in *D. suzukii* functionally. Gustatory perception of sugar in *Drosophila* depends on nine partially redundant chemoreceptors expressed in a complex combinatorial manner in ∼100 GRNs on different body parts and internally (reviewed in refs. ^1,28^). Targeting all nine receptors in *D. suzukii* is extremely challenging due to the lack of tools for genetic crosses. As an alternative, we thus generated in *D. suzukii* a pan-sugar-GRN Gal4 line homologous to the *DmelGr64af-Gal4* line previously shown to drive Gal4 expression in almost every sugar GRN ^29–31^. Fluorescent UAS reporters show expression of our *DsuzGr64af-Gal4* line in the main gustatory organs, the labellum and tarsi, in a highly reproducible pattern (**Fig. 4a-b**). Overall, fewer neurons are labelled in *D. suzukii* compared to *D. melanogaster* ^29–31^. We observe a reduction on the labellum (22 neurons vs ∼36 in *D. melanogaster*) and on specific tarsi of the foreleg (2 neurons vs 4 in the 4^th^ tarsal segment and 1 neuron vs 2 in the 3^rd^ and 2^nd^ tarsal segments). We confirmed this overall reduction with a second independent insertion line (not shown). Although we cannot completely rule out positional insertion effects of the Gal4 transgenes, this might reflect a general reduction of GRN numbers in *D. suzukii*, as has been reported for bitter GRNs ^21^. We also examined the axonal projections of GRNs labeled by *DsuzGr64af-Gal4* in the central brain and confirmed that they project exclusively to the Sub-Esophageal Zone (SEZ), with arborization patterns highly reminiscent of those of the equivalent line in *D. melanogaster* (**Supplementary Fig. 2**). To validate functionally our *DsuzGr64af-Gal4* line, we used it to silence these neurons in behavioral assays via a *UAS-Kir2*.*1* transgene ^32^ we generated in *D. suzukii*. Silencing sugar-GRNs with *Gr64af-Gal4* > *UAS-Kir2*.*1* significantly reduces the preference for sugar to a similar extent in both species in two-choice oviposition assays (**Fig. 4c**). In both species, sugar preference is not completely abolished however, suggesting an incomplete inhibition of sugar perception probably due to technical limitations (e.g., incomplete targeting of sugar-GRNs by the Gal4 transgenes and/or insufficient expression of the *UAS-Kir2*.*1* transgenes). In sum, our transgenic tools to manipulate sugar-GRNs in *D. suzukii* target similar sets of neurons and produce similar effects in behavioral assays as existing tools in *D. melanogaster*.

**Fig. 4.**
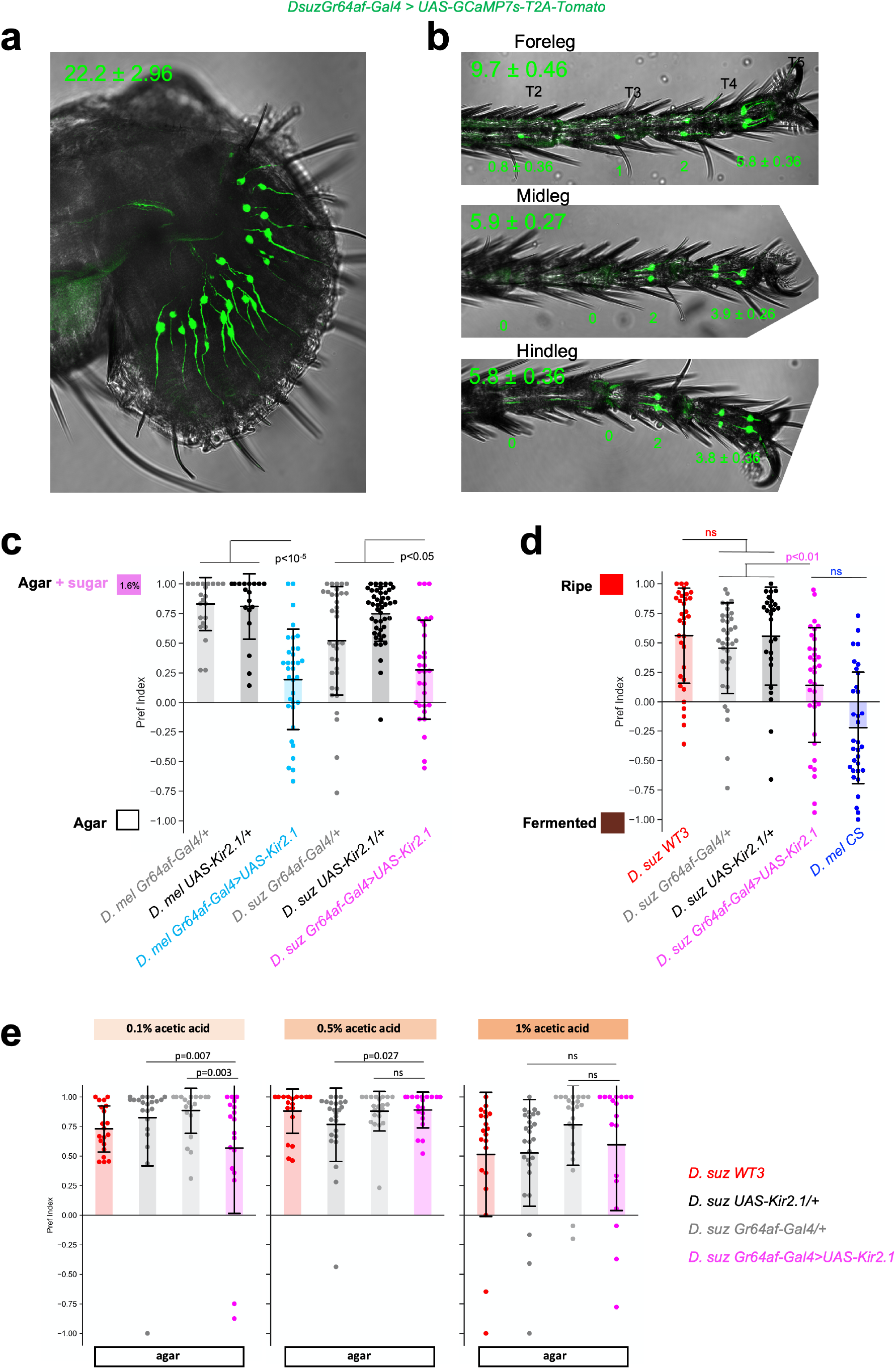
Sugar perception is required for ripe substrate preference in *D. suzukii*. **a, b** *DsuzGr64af-Gal4* expression in the proboscis (**a**) and legs (**b**) observed with a *UAS-GCaMP7s-T2A-Tomato* reporter. Numbers on top indicate the mean number of positive neurons ± standard deviation in the labellum and whole legs. In addition, the number of neurons in each tarsi are indicated in **b** (n=14 proboscises, forelegs, midlegs and hindlegs imaged). **c** Two-choice oviposition assay opposing 1.6% glucose+fructose to plain agar. Sugar-GRN inhibition via *UAS-Kir2*.*1* expression reduces - but does not completely abolish – the response to sugar to a similar extent in both species (n=25, 18, 34, 35, 46, 30). **d** Ripe vs fermented substrate oviposition choice assay. Sugar-GRN inhibition in *D. suzukii* interferes with oviposition decisions (n=33, 35, 26, 34, 33). **e** Two-choice oviposition assays opposing acetic acid at the indicated concentrations to plain agar, in the presence of 1% casein hydrolysate on both sides to stimulate egg-laying. Sugar-GRN inhibition does not abolish response to acetic acid (n=20, 25, 22, 18, 20, 25, 22, 19, 20, 25, 22, 19).

We then tested functionally the contribution of sugar perception to oviposition decisions on complex fruit substrates in *D. suzukii*. Remarkably, in two-choice assays opposing the ripe strawberry substrate to our fermented substrate, silencing sugar-GRNs significantly reduces *D. suzukii*’s preference for the ripe substrate (**Fig. 4d**). This manipulation brings *D. suzukii* close to indifference between the two substrates and closer to the *D. melanogaster* state. The preference is not completely abolished however, probably because of incomplete silencing and/ or the presence other instructive cues in the ripe substrate (see **Fig. 2c**).

Potential confounding effects with this experiment could arise from the fact that a subset of sugar-GRNs were shown in *D. melanogaster* to respond to other chemical cues in addition to sugar, including the fermentation product acetic acid ^33^ which can act as an attractant for oviposition ^33–35^. Silencing *D. suzukii* sugar-GRNs might thus interfere with oviposition decisions on the fermented substrate via affecting acetic acid perception. However, we ruled out this possibility as *DsuzGr64af-Gal4* > *UAS-Kir2*.*1* females display normal responses to acetic acid in two-choice oviposition assays at concentrations matching those of our fermented substrate (**Fig. 4e**). In conclusion, our results suggest that sugar perception in *D. suzukii* contributes significantly to oviposition decision-making on complex fruit substrates. Reducing sugar perception reduces the attractiveness of the ripe substrate over the fermented one, confirming that sugar is one of the major determinants of preference for ripe fruits in *D. suzukii*.

### Increasing sugar inputs in *D. melanogaster* reduces its preference for the fermented substrate

Together, our results suggest that evolutionary changes in neuronal circuits involved in encoding sugar value have participated in *D. suzukii*’s ecological shift of oviposition host. To further test this idea and as a proof of concept, we reasoned that artificially increasing the value of sugar in *D. melanogaster* should produce a similar shift in oviposition substrate preference. We thus decided to increase the sensitivity of sugar-GRNs in *D. melanogaster* to enhance their output on CNS oviposition decision-making circuits and thus increase the weight sugar exerts on the decision process.

To that end, we expressed the voltage-gated bacterial sodium channel NaChBac ^36^ in *D. melanogaster* sugar-GRNs with *Gr64af-Gal4* to make them hypersensitive. We first verified that this manipulation increases the behavioral response to sugar by performing egg-laying stimulation assays with glucose. As expected, raising sugar-GRN sensitivity dramatically increases the egg-laying rate on glucose compared to controls at all concentrations tested, in a manner resembling wild type *D. suzukii* behavior (**Fig. 5a**, raw data in **Supplementary Fig. 3**). *DmelGr64af-Gal4 > UAS-NaChBac* flies still respond to sugar in a dose-dependent manner, therefore their egg-laying response is amplified but their capacity to measure sugar concentration is not abolished. These observations are consistent with our earlier findings that the sugar concentration perceived by sensory neurons is directly translated into a quantitative oviposition response (**Fig. 3**). We then tested whether raising the perceived value of sugar can modify the preference in two-choice assays involving complex fruit substrates. The preference of *DmelGr64af-Gal4 > UAS-NaChBac* flies for the fermented substrate over the ripe is significantly reduced compared to genetic controls (**Fig. 5b**). Therefore, increasing sugar value shifts *D. melanogaster*’s preference towards ripe fruits, supporting the idea that an evolutionary change in sugar valuation could have contributed to *D. suzukii*’s behavioral shift. As observed when silencing sugar-GRNs in *D. suzukii*, the preference is not completely reversed and *DmelGr64af-Gal4 > UAS-NaChBac* flies still significantly prefer the fermented substrate. This could be due to the technical limitations in expression levels/patterns discussed previously and/ or could highlight the fact that other neuronal modifications are required to fully recapitulate the evolutionary transition that *D. suzukii* has undergone.

**Fig. 5.**
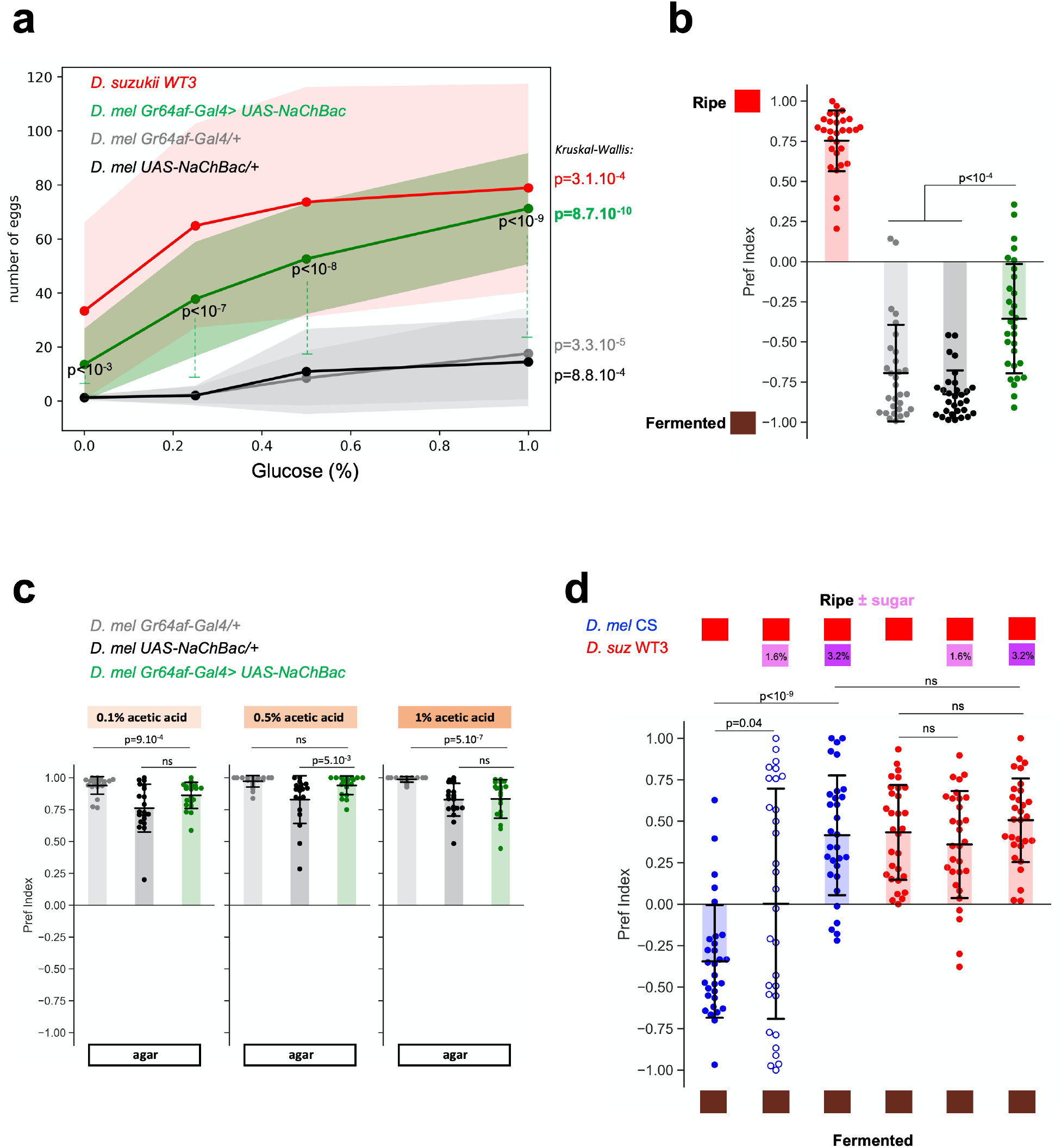
Increasing sugar inputs on the CNS in *D. melanogaster* biases oviposition substrate preference. **a** Oviposition stimulation assays (i.e.: no-choice) on increasing concentrations of glucose. Hypersensitizing *D. melanogaster* sugar-GRNs via *UAS-NaChBac* expression reproduces several characteristic features of *D. suzukii*’s oviposition responses to sugar; it increases the egg-laying rate on plain agar and all sugar concentrations (p-values shown on the graph indicate comparison to both parental controls) but does not abolish the capacity of the flies to adjust their oviposition response to sugar concentration (Kruskal-Wallis test, shown on the right of the graph) (n=30 for each condition). **b** Increasing sugar inputs in *D. melanogaster* significantly increases the value of the ripe substrate when opposed to a fermented substrate (n=30, 30, 30, 30). **c** Two-choice oviposition assays opposing acetic acid to plain agar, in the presence of 1% casein hydrolysate on both sides. Hypersensitizing sugar-GRNs does not significantly modify the preference for acetic acid (n=20, 20, 20, 20, 19, 20, 20, 20, 20). **d** Increasing the sugar concentration of the ripe substrate increases its value relative to the fermented substrate for *D. melanogaster*. One or two doses of sugar (1.6% or 3.2% glucose+fructose, indicated by pink boxes on top) were added to the ripe substrate (already containing 1.6% glucose+fructose) in two-choice assays against the fermented substrate. *D. melanogaster*’s preference for fermented is cancelled out when adding one dose and reversed to preference for the sweetened ripe substrate when adding two doses (n=30 for each condition).

As noted previously, manipulating sugar-GRN sensitivity could in theory interfere with oviposition decisions via modifying acetic acid perception since a subset of sugar-GRNs respond this compound. However, the two following experiments suggest otherwise. First, *DmelGr64af-Gal4 > UAS-NaChBac* flies showed normal responses to acetic acid in two-choice oviposition assays (**Fig. 5c**). In the second experiment, we specifically raised the perceived value of sugar without changing that of acetic acid: we used wild type flies and artificially raised the sugar concentration of the ripe substrate in a two-choice assay opposing it to the fermented substrate. Not surprisingly, adding sugar to the ripe substrate did not change the preference of wild-type *D. suzukii* for ripe (**Fig. 5d**). In sharp contrast, this manipulation shifted the preference of wild-type *D. melanogaster* towards the ripe substrate and even reversed its preference when using a high sugar concentration (i.e.: final conc. 3 times that of a regular ripe substrate; **Fig. 5d**). These experiments thus demonstrate that increasing the weight of sugar can bias *D. melanogaster*’s preference towards ripe fruit, supporting the idea that this is the main force that shifts the behavior of *DmelGr64af-Gal4 > UAS-NaChBac* flies in the ripe vs fermented choice assay. Remarkably, *D. melanogaster*’s preference for the high-sugar ripe substrate is indistinguishable from that of *D. suzukii* on the regular ripe substrate. Thus, raising the perceived sugar concentration appears sufficient to convert *D. melanogaster* into a *D. suzukii*-like state. In addition, these results are consistent with our previous observation that wild type *D. melanogaster* give high value to sugar only when present at a high concentration (**Fig. 2f**).

### Increased sugar valuation in *D. suzukii* is likely encoded by neuronal sensitivity changes at multiple levels of the circuit

Taken together, our results suggest an evolutionary scenario where changes in the neuronal circuits encoding sugar value have contributed to *D. suzukii*’s behavioral shift. These changes could have occurred directly at the level of the sensory neurons and/or in downstream circuits. A number of molecular and cellular mechanisms can be envisaged, starting with a change in the excitability of the sensory neurons, as suggested by our results with artificial manipulation of sugar-GRN sensitivity with NaChBac in *D. melanogaster*. As a first step in localizing the neuronal substrate of inter-species divergent sugar valuation, we decided to test this idea by comparing the excitability of sugar-GRNs between species using *in vivo* calcium imaging. We generated a *UAS-GCaMP7s* transgenic line in *D. suzukii* and monitored calcium responses in both species in the sugar-GRN synaptic terminals in the SEZ upon stimulation of the labellum with a range of glucose concentrations. Sugar stimulation of the labellum was shown to be sufficient to elicit egg-laying in *D. melanogaster* ^37^.

In both species, sugar-GRNs respond to glucose in a dose-dependent manner. The responses are detectable starting at 1% glucose but become significantly different from water controls starting at 2.5% glucose (**Fig. 6a-c**). Thus, we were unable to detect responses in *D. suzukii* at lower concentrations than in *D. melanogaster*, suggesting that increased sugar valuation is not simply due to increased sensitivity of the sensory neurons. In contrast, we did observe more important inter-species differences at high glucose concentrations. The magnitudes of the responses are greater for *D. melanogaster* at 5% and 10% glucose compared to *D. suzukii*, suggesting a higher dynamic range in *D. melanogaster* (highest ΔF/F_0_ ∼36% on average in *D. melanogaster* compared to ∼15% in *D. suzukii*, upon 5% and 10% glucose stimulation, respectively). In addition, we also observed differences in the temporal dynamics of the calcium responses: we consistently observed a rapid attenuation of the responses in *D. suzukii* after the ΔF/F_0_ peak which is almost absent in *D. melanogaster*, especially striking at high glucose concentrations (5-10% glucose; **Fig. 6a**). This suggests that signaling of sugar-GRNs to downstream circuits is sustained at higher levels and over longer periods of time in *D. melanogaster* at these concentrations.

**Fig. 6.**
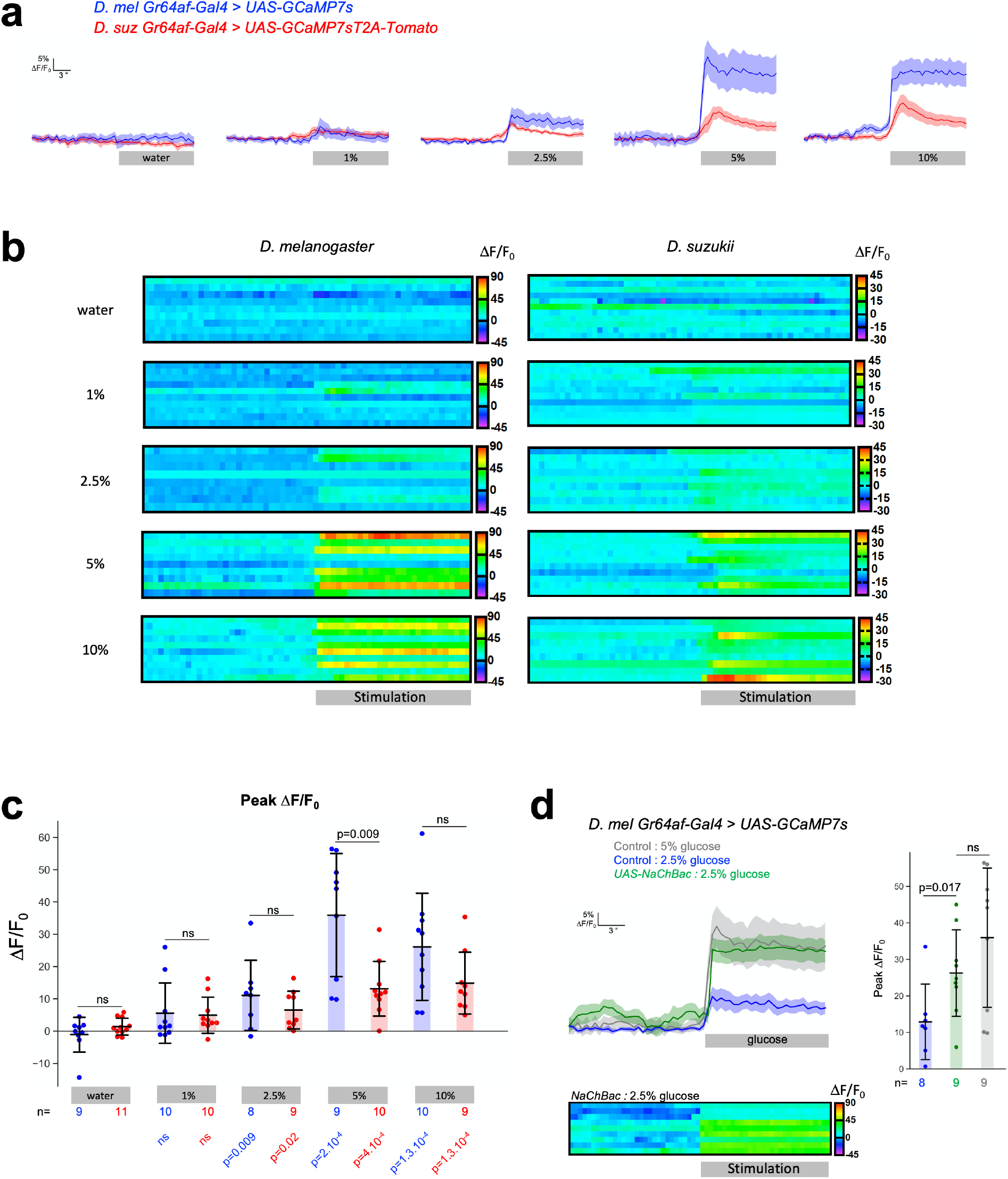
*D. melanogaster* behavioral tuning to high sugar concentrations correlates with higher calcium response in sugar-GRNs compared to *D. suzukii*. **a, b, c** GCaMP7s responses in sugar-GRNs upon stimulation of the labellum with increasing concentrations of glucose. **a** Average ΔF/F_0_ traces +/-s.e.m (shaded areas). The period of sugar stimulation is represented by the grey rectangles. **b** Individual traces represented as heat maps (note the different scales for *D. melanoagster* and *D. suzukii*; on the right). **c** Distributions of peak ΔF/F_0_ for each condition (shaded area: mean, error bars: standard deviation). Calcium response magnitudes are higher for *D. melanogaster* at high glucose concentrations, statistically significant for 5% glucose (Mann-Whitney U test). P-values in color below the graph indicate statistical comparisons to water controls and numbers above the p-values indicate sample sizes. **c** NaChBac expression in *D. melanogaster* sugar-GRNs significantly increases calcium responses to 2.5% glucose, up to levels similar to 5% glucose stimulation in controls. Average ΔF/F_0_ (top left), individual traces (bottom left) and peak ΔF/F_0_ (top right).

Since expressing NaChBac in *D. melanogaster* sugar-GRNs significantly enhanced its oviposition response to sugar (**Fig. 5**), we assessed how this manipulation affects the firing activity of the neurons. As expected, NaChBac expression significantly increased GCaMP responses to 2.5% glucose, up to levels comparable with 5% glucose stimulation in controls (**Fig. 6d**). This confirms that higher firing activity in sugar-GRNs induces a higher egg-laying rate and increases sugar weight in oviposition decisions.

Taken together, our results suggest that in *D. melanogaster*, a higher level of firing activity in the sugar-GRNs is required to trigger the oviposition program compared to *D. suzukii*. This suggests that the downstream CNS circuits are tuned to different levels of sugar-GRN signaling in the two species to promote oviposition starting at lower doses of sugar in *D. suzukii* than in *D. melanogaster*. Thus, increased sugar valuation in *D. suzukii* appears to be encoded by changes at multiple levels of the circuit.

## Discussion

We demonstrate that sugar has become a key chemosensory cue guiding oviposition decisions of *D. suzukii*. Glucose and fructose, abundant in ripe fruits, are a major source of nutrition for insects and can act as attractants for *Drosophila* oviposition in specific contexts ^26,27^. We find that the sugar content exerts a more important effect on *D. suzukii* oviposition decisions compared to *D. melanogaster* in several contexts and that sugar perception is required for *D. suzukii*’s oviposition attraction to ripe fruits. Thus, a change in the relative value of sugar - relative to other compounds including fermentation products – is responsible in part for *D. suzukii*’s oviposition preference shift.

The greater receptivity of *D. suzukii* to sugar in oviposition stimulation assays raises an apparent paradox: if this species readily accepts low-sugar substrates for egg-laying, it should not rely on the sugar concentration to discriminate high-sugar ripe strawberry from low-sugar fermented substrates in choice assays since both would be acceptable. However, we found that *D. suzukii* is clearly able to discriminate these sugar concentrations and is less stimulated by the sugar concentration of a fermented compared to a ripe substrate. Thus *D. suzukii* can in principle use sugar concentration as an indicator of substrate quality during oviposition decisions. Conversely, the lower receptivity of *D. melanogaster* to sugar should induce a preference for the high-sugar ripe substrate in choice assays if sugar was an important cue. However, we found evidence that sugar is unimportant to *D. melanogaster* as long as fermentation products are present and only becomes important when it is the sole cue present, or when present at high concentrations or perceived as such, as we observed when adding two doses of extra sugar to the ripe substrate (**Fig. 5d**) or raising sugar-GRN excitability with NaChBac (**Fig. 5b**). Taken together, these observations confirm the idea that the sugar relative value is highly context-dependent, and that sugar is valued very differently in the two species for choosing an oviposition site.

We explored several scenarios for the mechanisms underlying increased sugar valuation in *D. suzukii* at the level of the Peripheral Nervous System (PNS). First, an increase in the number of sugar-GRNs could result in stronger inputs onto the Central Nervous System (CNS), resulting in increasing the value of sugar. However, our *Gr64af-Gal4* reporter analysis suggests otherwise since we detected fewer sugar-GRNs in *D. suzukii* than *D. melanogaster*. While we cannot exclude positional effects of the transgenes, our functional analyses suggest that the reporters target a similar proportion of sugar-GRNs in the two species. Thus *D. suzukii* appears to harbor less sugar-GRNs than *D. melanogaster* as reported also for bitter-GRNs ^21^. We cannot exclude that more subtle differences in the numbers of functionally-distinct sugar-GRN subtypes exist between species. Indeed, different sub-populations of sugar-GRNs in *D. melanogaster* exert opposite effects on oviposition behavior, with GRNs on the legs inhibiting oviposition while GRNs on the proboscis promoting oviposition in a specific context ^37^. Interestingly, we observed a reduction of sugar-GRNs in tarsal segments 2, 3 and 4 of the *D. suzukii* foreleg compared to *D. melanogaster*, raising the possibility that disappearance of specific neuronal subtypes could have relieved inhibitory sugar inputs onto CNS oviposition circuits in this species. Another possible mechanism for increased sugar valuation in *D. suzukii* would be an increased sensitivity of sugar-GRNs. We explored this idea via calcium imaging experiments but detected in fact higher GCaMP responses (in magnitude and duration) in *D. melanogaster* compared to *D. suzukii* upon stimulation at high glucose concentrations. This suggests that in this regime, sugar inputs onto the CNS are stronger in *D. melanogaster*. These observations can be reconciled with our behavioral data if we hypothesize that the evolutionary difference between species also includes changes in the tuning of the CNS circuits triggering oviposition in response to sugar inputs (**Fig. 7a**). More precisely, we hypothesize that the CNS circuits are more sensitive to PNS sugar inputs in *D. suzukii* than in *D. melanogaster*, concomitant with divergent responses to fermentation compounds as well. Thus, in *D. suzukii*, the CNS circuits downstream of sugar perception would require low inputs to trigger oviposition, consistent with the enhanced behavioral response to low sugar concentrations and the flatter dynamic range of the sensory neurons. In contrast, in *D. melanogaster*, the CNS circuits would require higher sugar inputs to trigger oviposition. This is exemplified by the higher GCaMP response magnitude and duration in the PNS occurring at the sugar concentrations required to elicit the maximal oviposition response in *D. melanogaster*. This model is consistent with the fact that artificially raising sugar inputs in *D. melanogaster* via NaChBac enhances its behavioral response to sugar and increases ripe substrate preference (**Fig. 5a-b**), as does raising the sugar concentration of the ripe substrate (**Fig. 5d**). These data are also consistent with the fact that *D. melanogaster* is a better discriminator of sugar concentration than *D. suzukii*, as we did observe in competition experiments opposing two sugar concentrations (**Fig. 2h**).

**Fig. 7.**
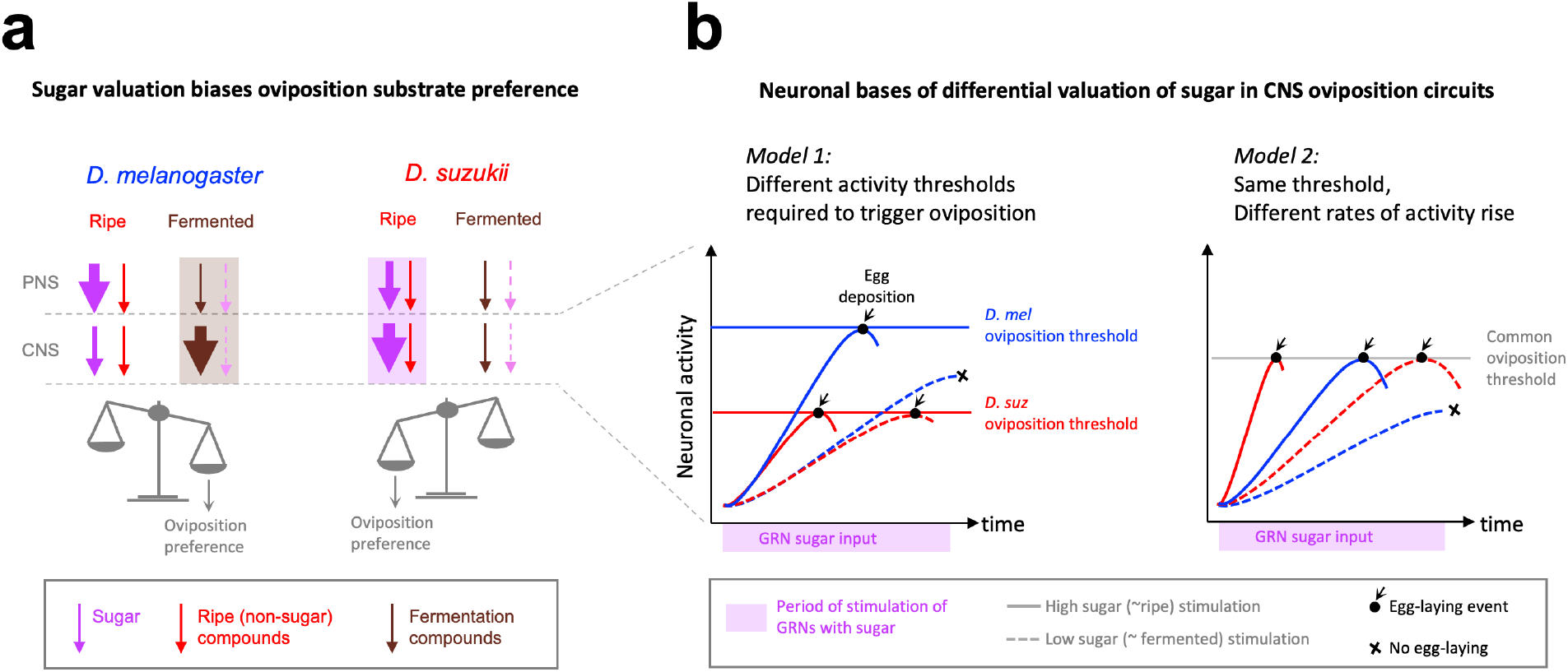
Model – Interspecies differences in sugar valuation underlie divergent oviposition substrate preferences. **a** schematic representing the integration of chemosensory cues in both species in a ripe vs fermented substrate choice assay. Neuronal activity in sensory neurons (PNS; top row) and CNS oviposition circuits (bottom row) is represented by arrows (thickness represents activity level) associated to specific chemosensory cues: sugar (purple), non-sugar cues present in ripe fruits (red) and fermentation compounds (brown). In *D. melanogaster*, sugar from the ripe substrate elicits high responses in the PNS but low responses in CNS, resulting in a relatively low weight on the decision. In contrast, fermentation compounds are assigned higher value in the PNS and/or CNS (shown here in CNS), overall biasing the preference towards the fermented substrate (brown shaded rectangle). In *D. suzukii*, sugar elicits lower responses in the PNS but these are amplified in the CNS, thereby assigning high value, while fermentation cues elicit low responses, resulting overall in a preference for the ripe substrate (pink shaded rectangle). **b** Hypothetical neuronal mechanisms underlying interspecies differential sugar valuation in the CNS. Neuronal activity (y-axis) rises over time (x-axis) in oviDN neurons upon stimulation with sugar from a ripe (solid lines) or fermented substrate (dashed lines) until reaching the oviposition threshold (“egg deposition”, black dot and arrow). Failure to reach this threshold aborts egg-laying (black cross). Two scenarios relying on differences in physiological properties of oviposition circuit neurons (oviDN and/or upstream neurons) could provide a mechanistic basis for divergent sugar valuation between species under these conditions: (*Model 1*) the oviDN activity threshold required to trigger oviposition is lower in *D. suzukii* (red) than in *D. melanogaster* (blue). (*Model 2*) the threshold is the same in both species but the slope of activity rise in response to sugar is higher in *D. suzukii*. In both scenarios, lower signaling from the PNS would be required in *D. suzukii* to reach the CNS activity threshold sufficient to trigger oviposition without affecting its ability to discriminate ripe from fermented substrates. This would effectively increase the probability of *D. suzukii* laying an egg in response to sugar compared to *D. melanogaster*.

Which CNS neurons and neurophysiological differences could be involved in the species divergence of sugar valuation? The CNS circuits downstream of sugar gustatory perception and controlling oviposition remain poorly characterized in *D. melanogaster*. Connectivity analyses, calcium imaging studies and functional approaches have identified several populations of direct and indirect sugar-GRN targets regulating mostly feeding behaviors ^38–47^. In addition, three CNS neuronal populations have been implicated in regulating oviposition including in response to sugar: the TPN2 neurons ^37^, dopaminergic neurons ^46,48^ and the oviIN-oviEN-oviDN circuit ^49^. These neurons thus represent interesting candidates to investigate the neuronal bases of species differential sugar valuation. Recent data suggests that the oviDN neurons act as a temporal accumulator of sensory and internal drive for oviposition and enact the decision to lay an egg upon reaching a specific threshold of activity. The rate at which activity rises in oviDN neurons appears to encode substrate attractivity for oviposition ^50^. This raises the exciting hypothesis that inter-species divergent sugar valuation could arise from physiological differences in oviposition circuits affecting the rate of activity rising or threshold height required in oviDN neurons to trigger oviposition in response to sugar stimulation (**Fig. 7b**). Future work should thus aim at identifying the functional homologs of the different groups of neurons composing the oviposition circuit in the *D. suzukii* brain and assessing how they are regulated by or interact with oviposition cues including sugar. Functional imaging of large brain areas ^47^ will probably also shed light on as yet unidentified neuronal pathways downstream of sugar perception which might harbor inter-species divergence in sugar valuation processes.

## Methods

### *Drosophila* stocks

Flies were reared on home-made Nutrifly medium or standard cornmeal medium when indicated, at 23°C, 60% relative humidity in a 12:12hr light cycle. Wild type *D. melanogaster* were Canton S, Canton S SNP iso2 (BL6365), OreR, Harwich (BL4264) and iso1 (isofemale line from India, B. Prud’homme). Wild type *D. suzukii* were WT3 ^51^, an isofemale line from Japan (JAP) and an isofemale line from France (AM). Other wild type species in **Fig. 1c**: (*bia*) 3 isofemale *D. biarmipes* lines from Bangalore, India (b1, b2, b3); (*erec*) *D. erecta* #223; (*sec*) *D. sechellia* A1 (14021-0248.28) and A2 (14021-0248.07); (*sim*) *D. simulans md221#*343 (P. andolfatto) and *D. simulans Vincennes#347*, obtained from V. Courtier-Orgogozo; (*tak*) *D. takahashii*; (*eug*) *D. eugracilis*; (*sant*) *D. santomea;* (*teiss*) *D. teissieri;* (*pseu*) *D. pseudoobscura;* and (*will*) *D. willistoni*.

*D. melanogaster* transgenic lines: *Gr64af-Gal4* (BL57668), *UAS-Cd4tdTomato* (BL35841), *UAS-Kir2*.*1* ^32^, *UAS-NaChBac* ^36^, *UAS-GCaMP7s* (BL80905).

### Transgenic *D. suzukii* lines

*-DsuzGr64af-Gal4*^*3xp3GFP*^ : a 10483bp fragment spanning the *Gr64a* 5’ to *Gr64f* 5’ region from the *D. suzukii* genome was synthesized by Genewiz and cloned into our PiggyBac transformation vector ^20^. This fragment is orthologous to the 10kb fragment used in the *DmelGr64af-Gal4* line ^29^.

-The *UAS-Cd4tdTomato*^*3xp3RFP*^ line was described in ref. ^20^.

-The *UAS-GCaMP7s-T2A-Tomato*^*3xp3RFP*^ line was generated via PiggyBac transformation of a vector obtained from D. Stern.

*-UAS-Kir2*.*1*^*aUP2-3xp3RFP*^ line: To facilitate future transgenic approaches, we first generated an attP landing site in *D. suzukii* via CRISPR-mediated homologous recombination, following our CRISPR protocol described in ref. ^20^. To design sgRNA oligos, we amplified and sequenced the *D. suzukii* genomic region orthologous to the *D. melanogaster* attP2 locus on chromosome 3. We targeted the following N18GG sequence: AGTTGTTTATAAACTAGGTGG with the sgRNA-F oligo (all primers sequences given below), coupled with the generic oligo sgRNA-R and co-injected with the sgRNA-attP2 oligo to insert the attP site. We screened for transformants by PCR and sequencing, using suzssODNattP2-F1 and suzssODNattP2-R primers for PCR and suzssODNattP2-F2 and suzssODNattP2-R primers for sequencing.

We PCR-amplified the Kir2.1 sequence from the *D. melanogaster* transgenic line using Kir2.1_Fw1 and Kir2.1_Rv1 primers and cloned it by in-fusion into a pBac-attB-UAS vector created from the PiggyBac vector used in ^20^. We then inserted the *UAS-Kir2*.*1* transgene at the *D. suzukii* attP2 site via the PhiC31 integrase system following the protocol described in ^52^: phiC31 mRNA was produced with the mMessage mMachine kit (Ambion, Austin, TX) with 1 μg of *Bam*HI-linearized pET11phiC31poly(A) plasmid template. phiC31 integrase-capped mRNA was injected in embryos at 1000 ng/μl, together with 200 ng/μl of UAS-Kir2.1 plasmid.

### Primers

sgRNA-F:

GAAATTAATACGACTCACTATAGGAGTTGTTTATAAACTAGGGTTTTAGAGCTAGAAATAGC

sgRNA-R:

AAAAGCACCGACTCGGTGCCACTTTTTCAAGTTGATAACGGACTAGCCTTATTTTAACTTGCTATTTCTAGCTCTAAAAC.

sgRNA-attP2:

CGGTGGTACTTGTGGGTATATGAAGCCGATTGGAGGGTTCCGACTGCACCCGGCGAGAGTTGTTTATAAACTAGTAGTGCCCCAACTGGGGTAACCTTTGAGTTCTCTCAGTTGGGGGCGTAGAATTCAGGTGGAGAGTGGCGATCATATGGACAAGCTTGCCATTCGCCAAAGCTTTCTAATACCAGTGTTAGTTAGCT

suzssODNattP2-F1 : CTTCAGGTAACCGGTTGTGG

suzssODNattP2-R: ATGCTGCCAAAGCAGTGTCT

suzssODNattP2-F2 : GGGGTACGAGGTGTTTTTAAG

Kir2.1_Fw1 : atgggcagtgtgcgaaccaaccgc

Kir2.1_Rv1 : tcatatctccgactctcgccgtaagg

### Strawberry fermentation

Frozen organic non-sweetened strawberry purée was purchased from Sicoly (https://www.sicoly.fr/). The purée was diluted 1:2 in milliQ water and 2g glucose / 100ml was added at the start of fermentation. For each fermentation, fresh overnight liquid cultures of *Saccharomyces cerevisiae* (gift from A. Michelot) and *Acetobacter pomorum* (gift from J. Royet) were grown at 30°C in standard media (YPD and MRS, respectively) and inoculated at a final concentration of 10^6^ cells/ml of diluted strawberry purée. Fermentation was performed for 72hrs in large Erlenmeyer flasks in a shaking incubator at 30°C. Fermented purée was stored at 4°C and used over a few days/weeks.

Glucose, fructose, acetic acid and ethanol dosages of fermented and unfermented purées were performed with kits from Megazyme (K-FRUGL, K-ACET, K-ETOH).

### Chemicals

Glucose (sigma G7021), fructose (sigma F0127), sucrose (sigma S1888), acetic acid (Carlo Erba 524520), grape juice (local casino store), casein hydrolysate (sigma 22090), agar (vwr 20768.361).

### Behavioral experiments

#### Experimental setups

experiments were performed in custom-built setups - “large” and “small” arenas - adapted from ref. ^53^, except the sugar dose-response experiments from **Fig. 3** that were performed in the experimental setup described in ref. ^20^; see below. The large and small setups consist of sheets of drilled plexiglass assembled to form large (6×4×1.8 cm) or small (3×2×1.8 cm) chambers each containing at their bottom two 0.8 cm-wide strips of oviposition substrate on opposite ends of the chamber (see schematics in **Fig. 1-2**). Three females were introduced per chamber. We found that oviposition experiments involving volatile compounds (i.e.: fruit purées, acetic acid) require the large setups which have increased air circulation compared to the small setups. Therefore, the small setups were only used for experiments involving non-volatile compounds (i.e.: sugar-alone).

Dose-response experiments from **Fig. 3** were performed in 12×6×4 cm chambers with ten females per chamber and one 4 cm-diameter Petri dish containing the oviposition substrate.

#### Oviposi9on assays

flies were collected between 0-3 days after eclosion, aged in fresh food vials for 7-9 days, sorted on CO_2_ just before the experiments, placed in behavioral chambers and left to lay eggs in the chambers for 20-24hrs in the dark. Eggs were counted manually and a preference index (PI) was calculated for each replica (for 2-choice experiments) using the following formula: PI = (# eggs on substrate 1 - # eggs substrate 2)/total # eggs on both substrates. Chambers with less than 5 eggs total were excluded from PI calculation. Oviposition substrates were prepared by mixing the indicated compounds with a solution of agar boiled separately to achieve a final concentration of 0.5% (w/v) agar. Fruit purées/juice were diluted to 30% (w/v) final concentration. For acetic acid oviposition preference assays, a source of protein (1% casein hydrolysate) was added in both substrates to stimulate egg-laying since acetic acid alone does not stimulate.

#### Data analysis and sta9s9cs

analyses, statistics and graphs were performed using custom-written scripts in Python. For two-choice experiments, the preference of each genotype/ condition for one of the two substrates was evaluated using a Mann-Whitney paired test comparing the # of eggs on substrate 1 vs # of eggs on substrate 2. p-values<0.05 indicate a significant preference for one of the two substrates (filled circles on the graphs), p-values>0.05 indicate no significant preference for either substrate (open circles). To determine if different genotypes/conditions showed different preferences in a given choice assay, their distributions of preference indexes were compared by a Mann-Whitney U-test (p-values indicated on the graphs). Distributions of total # of eggs laid were compared by Mann-Whitney U-test as well. Dose-response experiments were analyzed using a Kruskal-Wallis H-test to determine if each genotype/condition was significantly induced to lay eggs by the indicated sugar. The Effective Dose 50 was calculated as ED50 = (maximal rate - basal rate)/2.

### Microscopy

*Dsuz-Gr64af-Gal4* expression was examined in female live legs and proboscises using tomato expression from our *UAS-GCaMP7s-T2A-Tomato* reporter line. Legs and proboscises were mounted in 1xPBS.

*Dsuz-* and *Dmel-Gr64af-Gal4* projection patterns in the brain (SEZ) were examined by immunostaining against tomato (using *UAS-Cd4tdTomato* reporters). Immunostaining was performed as described in ref. ^20^ with the following antibodies: rabbit anti-RFP (Rockland, used at 1:1000), mouse nc82 (Hybridoma bank, used at 1:20), secondaries were anti-rabbit alexa 488 and anti-mouse alexa 647 (Rockland, used at 1:200). Brains were mounted in SlowFade medium (ThermoFisher). Images were acquired with a zeiss LSM780 confocal.

### Calcium imaging

*in vivo* calcium imaging was performed on 5-7-day-old mated females raised under the same conditions as behavioral experiments but starved for 24 hrs prior to the experiments. Flies were anesthetized on ice for 1 hr, suspended by the neck on a plexiglass block (2 × 2 × 2.5 cm) with the proboscis facing the center of the block and immobilized using an insect pin (0.1 mm diameter) placed on the neck and fixed on the block with beeswax (Deiberit 502, Siladent, 209212). To prevent movement, the head was then glued on the block with a drop of resin (Gum rosin, Sigma-Aldrich −60895-, dissolved in ethanol at 70 %) such that the anterior part of the head would be facing the microscope objective. Flies were then placed in a humidified box for 1 hr to allow the resin to harden. A plastic coverslip with a small hole (diameter ∼ distance between the two eyes) was placed on top of the head, fixed on the block with beeswax and then sealed on the cuticle with two-component silicon (Kwik-Sil, World Precision Instruments), leaving the proboscis exposed to air below the coverslip. A drop of Ringer’s saline (130 mM NaCl, 5 mM KCl, 2 mM MgCl_2_, 2 mM CaCl_2_, 36 mM sucrose, 5 mM HEPES, pH 7.3) was placed on the head and the cuticle spanning the antennae area was then removed. To allow visual access to the anterior-ventral part of the SEZ, the trachea and fat body were removed and the gut was cut without damaging the brain or taste nerves. The exposed brain was then rinsed twice with Ringer’s saline.

Imaging was performed with a Leica DM600B microscope with a 25x water objective. GCaMP7s fluorescence was excited using a Lumencor diode light source at 482 nm ± 25 and collected through a 505-530 nm band-pass filter. Images were acquired every 500 ms using a Hamamatsu/HPF-ORCA Flash 4.0 camera and processed using Leica MM AF 2.2.9.

Stimulation was performed by applying 140 μL of glucose (sigma) diluted in water on the proboscis. Each experiment consisted of a recording of 100 images before stimulation and 100 images after stimulation. Graphical representations show 30 images before and 30 images after stimulation.

Data processing was performed as described in ref. ^54^. For each recording, ROIs were drawn manually on the left and right side of the SEZ in FIJI (https://fiji.sc/) and signal intensity quantified. These data were then inspected manually to exclude recordings showing clear signs of drift and to select the side with the least fluctuations in the response. The initial intensity F_0_ was calculated over frames 1-10 and ΔF/F_0_ expressed as a %. Peak ΔF/F_0_ was calculated as the average ΔF/F_0_ over 4 frames around the peak / average ΔF/F_0_ over 4 frames immediately before the peak. Statistical difference in peak ΔF/F_0_ across species was assessed by Mann-Whitney U test.

## Acknowledgments

We are grateful to the Bloomington Drosophila Stock Center, L. Kurz, V. Courtier-Orgogozo, R. Benton, T. Auer, J. Blau and P. Andolfatto for fly stocks; D. Stern for the UAS-GCaMP7s-T2A-Tomato plasmid; Flybase for information support; J. Royet and A. Michelot for strains of *A. pomorum* and *S. cerevisae*, respectively.

## Author contributions

MC and BP conceived the project and designed experiments. MC, BC, SQ and ST performed behavioral experiments. GM performed calcium imaging experiments. MBG analyzed calcium imaging experiments. CM performed molecular biology and transgenesis, with BC. BD designed and built the behavioral setups. MC analyzed experimental results and wrote the manuscript with BP. BC, GM, MBG and YG edited and commented the manuscript.

## References

1. Joseph, R. M. & Carlson, J. R. Drosophila Chemoreceptors: A Molecular Interface Between the Chemical World and the Brain. Trends Genet (2015) doi:10.1016/j.tig.2015.09.005.

2. Auer, T. O. et al. Olfactory receptor and circuit evolution promote host specialization. Nature 579, 402–408 (2020).

3. Dekker, T., Ibba, I., Siju, K. P., Stensmyr, M. C. & Hansson, B. S. Olfactory shifts parallel superspecialism for toxic fruit in Drosophila melanogaster sibling, D. sechellia. Curr Biol 16, 101–9 (2006).

4. Matsuo, T., Sugaya, S., Yasukawa, J., Aigaki, T. & Fuyama, Y. Odorant-binding proteins OBP57d and OBP57e affect taste perception and host-plant preference in Drosophila sechellia. PLoS Biol 5, e118 (2007).

5. Prieto-Godino, L. L. et al. Evolution of Acid-Sensing Olfactory Circuits in Drosophilids. Neuron 93, 661–676 e6 (2017).

6. Linz, J. et al. Host plant-driven sensory specialization in Drosophila erecta. Proc Biol Sci 280, 20130626 (2013).

7. Mansourian, S. et al. Wild African Drosophila melanogaster Are Seasonal Specialists on Marula Fruit. Curr. Biol. 28, 3960–3968.e3 (2018).

8. Wisotsky, Z., Medina, A., Freeman, E. & Dahanukar, A. Evolutionary differences in food preference rely on Gr64e, a receptor for glycerol. Nat Neurosci 14, 1534–41 (2011).

9. McBride, C. S. et al. Evolution of mosquito preference for humans linked to an odorant receptor. Nature 515, 222–7 (2014).

10. Wada-Katsumata, A., Silverman, J. & Schal, C. Changes in taste neurons support the emergence of an adaptive behavior in cockroaches. Science 340, 972–5 (2013).

11. Grosjean, Y. et al. An olfactory receptor for food-derived odours promotes male courtship in Drosophila. Nature 478, 236–40 (2011).

12. McGrath, P. T. et al. Parallel evolution of domesticated Caenorhabditis species targets pheromone receptor genes. Nature 477, 321–5 (2011).

13. de Bono, M. & Bargmann, C. I. Natural variation in a neuropeptide Y receptor homolog modifies social behavior and food response in C. elegans. Cell 94, 679–89 (1998).

14. Ding, Y., Berrocal, A., Morita, T., Longden, K. D. & Stern, D. L. Natural courtship song variation caused by an intronic retroelement in an ion channel gene. Nature 536, 329–32 (2016).

15. Seeholzer, L. F., Seppo, M., Stern, D. L. & Ruta, V. Evolution of a central neural circuit underlies Drosophila mate preferences. Nature 559, 564–569 (2018).

16. Asplen, M. K. et al. Invasion biology of spotted wing Drosophila (Drosophila suzukii): a global perspective and future priorities. J. Pest Sci. 88, 469–494 (2015).

17. Cini, A. A review of the invasion of Drosophila suzukii in Europe and a draft research agenda for integrated pest management. Bull. Insectology 65, 149–160 (2012).

18. Atallah, J., Teixeira, L., Salazar, R., Zaragoza, G. & Kopp, A. The making of a pest: the evolution of a fruit-penetrating ovipositor in Drosophila suzukii and related species. Proc. R. Soc. B Biol. Sci. 281, 20132840 (2014).

19. Green, J. E. et al. Evolution of Ovipositor Length in Drosophila suzukii Is Driven by Enhanced Cell Size Expansion and Anisotropic Tissue Reorganization. Curr. Biol. 29, 2075–2082.e6 (2019).

20. Karageorgi, M. et al. Evolution of Multiple Sensory Systems Drives Novel Egg-Laying Behavior in the Fruit Pest Drosophila suzukii. Curr Biol 27, 847–853 (2017).

21. Dweck, H. K., Talross, G. J., Wang, W. & Carlson, J. R. Evolutionary shifts in taste coding in the fruit pest Drosophila suzukii. eLife 10, e64317 (2021).

22. Durkin, S. M. et al. Behavioral and Genomic Sensory Adaptations Underlying the Pest Activity of Drosophila suzukii. Mol. Biol. Evol. 38, 2532–2546 (2021).

23. May, C. E. et al. High Dietary Sugar Reshapes Sweet Taste to Promote Feeding Behavior in Drosophila melanogaster. Cell Rep. 27, 1675–1685.e7 (2019).

24. Montero, T. M., Mollá, E. M., Esteban, R. M. & López-Andréu, F. J. Quality attributes of strawberry during ripening. Sci. Hortic. 65, 239–250 (1996).

25. USDA, strawberry content. https://fdc.nal.usda.gov/fdc-app.html#/food-details/2263887/nutrients. US Department of Agriculture https://fdc.nal.usda.gov/fdc-app.html#/food-details/2263887/nutrients.

26. Schwartz, N. U., Zhong, L., Bellemer, A. & Tracey, W. D. Egg laying decisions in Drosophila are consistent with foraging costs of larval progeny. PLoS One 7, e37910 (2012).

27. Yang, C. H., Belawat, P., Hafen, E., Jan, L. Y. & Jan, Y. N. Drosophila egg-laying site selection as a system to study simple decision-making processes. Science 319, 1679–83 (2008).

28. Freeman, E. G. & Dahanukar, A. Molecular neurobiology of Drosophila taste. Curr Opin Neurobiol 34, 140–8 (2015).

29. Dahanukar, A., Lei, Y. T., Kwon, J. Y. & Carlson, J. R. Two Gr genes underlie sugar reception in Drosophila. Neuron 56, 503–16 (2007).

30. Fujii, S. et al. Drosophila sugar receptors in sweet taste perception, olfaction, and internal nutrient sensing. Curr Biol 25, 621–627 (2015).

31. Ling, F., Dahanukar, A., Weiss, L. A., Kwon, J. Y. & Carlson, J. R. The molecular and cellular basis of taste coding in the legs of Drosophila. J Neurosci 34, 7148–64 (2014).

32. Baines, R. A., Uhler, J. P., Thompson, A., Sweeney, S. T. & Bate, M. Altered electrical properties in Drosophila neurons developing without synaptic transmission. J Neurosci 21, 1523–31 (2001).

33. Chen, H.-L., Stern, U. & Yang, C.-H. Molecular control limiting sensitivity of sweet taste neurons in Drosophila. Proc. Natl. Acad. Sci. 116, 20158–20168 (2019).

34. Chen, Y. & Amrein, H. Ionotropic Receptors Mediate Drosophila Oviposition Preference through Sour Gustatory Receptor Neurons. Curr Biol 27, 2741–2750 e4 (2017).

35. Joseph, R. M., Devineni, A. V., King, I. F. & Heberlein, U. Oviposition preference for and positional avoidance of acetic acid provide a model for competing behavioral drives in Drosophila. Proc Natl Acad Sci U A 106, 11352–7 (2009).

36. Nitabach, M. N. Electrical Hyperexcitation of Lateral Ventral Pacemaker Neurons Desynchronizes Downstream Circadian Oscillators in the Fly Circadian Circuit and Induces Multiple Behavioral Periods. J. Neurosci. 26, 479–489 (2006).

37. Chen, H.-L., Motevalli, D., Stern, U. & Yang, C.-H. A functional division of Drosophila sweet taste neurons that is value-based and task-specific. Proc. Natl. Acad. Sci. 119, e2110158119 (2022).

38. Miyazaki, T., Lin, T.-Y., Ito, K., Lee, C.-H. & Stopfer, M. A gustatory second-order neuron that connects sucrose-sensitive primary neurons and a distinct region of the gnathal ganglion in the Drosophila brain. J. Neurogenet. 29, 144–155 (2015).

39. Talay, M. et al. Transsynaptic Mapping of Second-Order Taste Neurons in Flies by trans-Tango. Neuron 96, 783–795.e4 (2017).

40. Flood, T. F. et al. A single pair of interneurons commands the Drosophila feeding motor program. Nature 499, 83–87 (2013).

41. Gordon, M. D. & Scott, K. Motor Control in a Drosophila Taste Circuit. Neuron 61, 373–384 (2009).

42. Kain, P. & Dahanukar, A. Secondary taste neurons that convey sweet taste and starvation in the Drosophila brain. Neuron 85, 819–32 (2015).

43. Kim, H., Kirkhart, C. & Scott, K. Long-range projection neurons in the taste circuit of Drosophila. eLife 6, e23386 (2017).

44. Liu, C. et al. A subset of dopamine neurons signals reward for odour memory in Drosophila. Nature 488, 512–516 (2012).

45. Yapici, N., Cohn, R., Schusterreiter, C., Ruta, V. & Vosshall, L. B. A Taste Circuit that Regulates Ingestion by Integrating Food and Hunger Signals. Cell 165, 715–729 (2016).

46. Vijayan, V. et al. An internal expectation guides Drosophila egg-laying decisions. http://biorxiv.org/lookup/doi/10.1101/2021.09.30.462671 (2021) doi:10.1101/2021.09.30.462671.

47. Harris, D. T., Kallman, B. R., Mullaney, B. C. & Scott, K. Representations of Taste Modality in the Drosophila Brain. Neuron 86, 1449–1460 (2015).

48. Yang, C. H., He, R. & Stern, U. Behavioral and circuit basis of sucrose rejection by Drosophila females in a simple decision-making task. J Neurosci 35, 1396–410 (2015).

49. Wang, F. et al. Neural circuitry linking mating and egg laying in Drosophila females. Nature 579, 101–105 (2020).

50. Vijayan, V. et al. A rise-to-threshold signal for a relative value deliberation. http://biorxiv.org/lookup/doi/10.1101/2021.09.23.461548 (2021) doi:10.1101/2021.09.23.461548.

51. Chiu, J. C. et al. Genome of Drosophila suzukii, the Spotted Wing Drosophila. G3 GenesGenomesGenetics 3, 2257–2271 (2013).

52. Groth, A. C. Construction of Transgenic Drosophila by Using the Site-Specific Integrase From Phage C31. Genetics 166, 1775–1782 (2004).

53. Gou, B., Zhu, E., He, R., Stern, U. & Yang, C. H. High Throughput Assay to Examine Egg-Laying Preferences of Individual Drosophila melanogaster. J Vis Exp e53716 (2016) doi:10.3791/53716.

54. Silbering, A. F., Bell, R., Galizia, C. G. & Benton, R. Calcium Imaging of Odor-evoked Responses in the Drosophila Antennal Lobe. J. Vis. Exp. 2976 (2012) doi:10.3791/2976.

