## Supplementary figures for "Evolutionary divergence in sugar valuation shifts *Drosophila suzukii* oviposition choice towards ripe fruit"

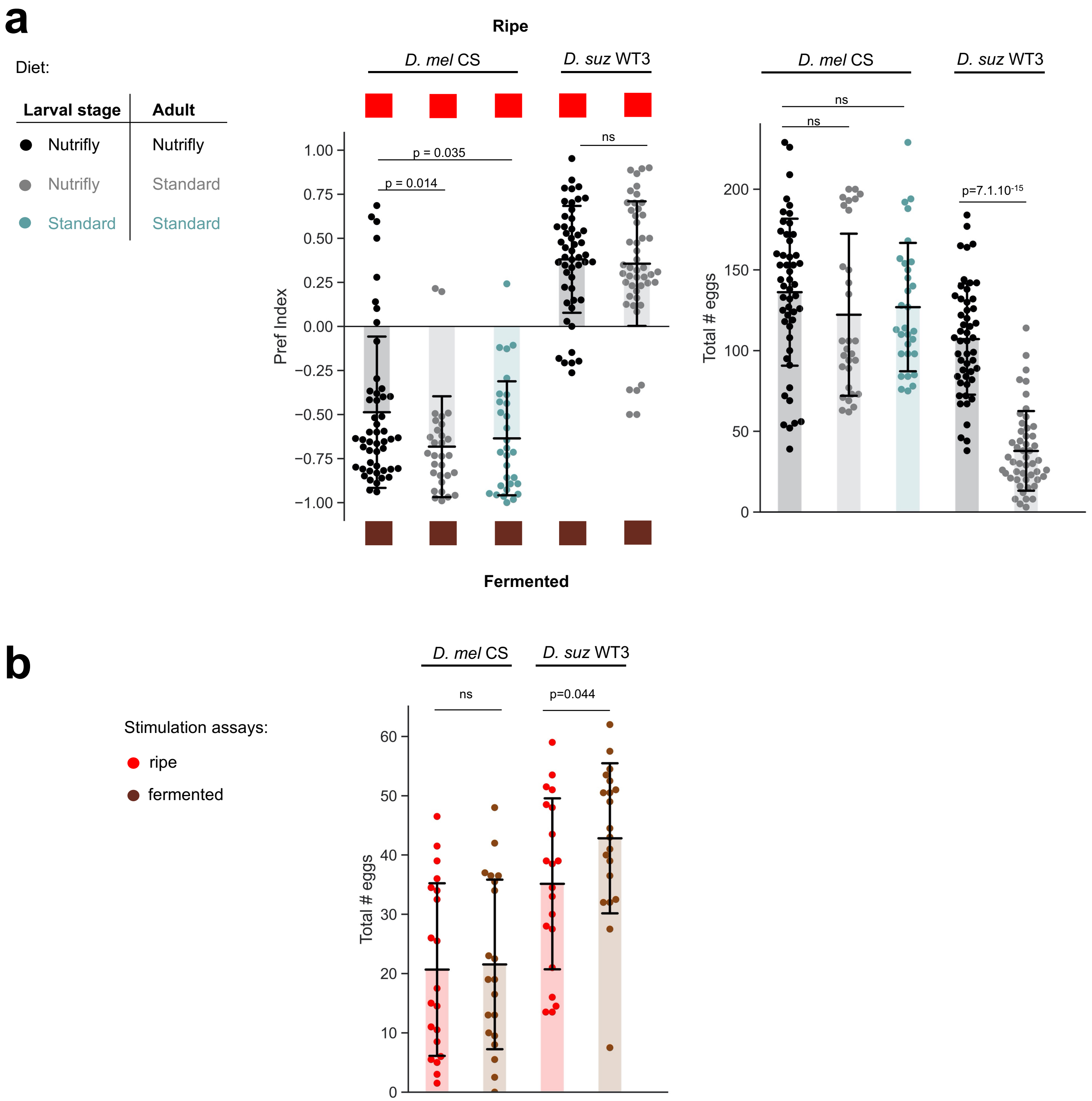


**Supplementary Fig. 1.** Control experiments with the ripe and fermented substrates.

**a** The diet does not significantly modify species oviposition preferences for ripe vs fermented substrates. The two species were allowed to develop on Nutrifly medium during the larval stages and either kept on this medium during adulthood (black) or switched to standard cornmeal medium at eclosion (grey). *D. melanogaster* were also grown directly on standard medium during larval stages (light blue), but we did not manage to grow *D. suzukii* under these conditions. Left: the oviposition preference is independent of diet, right: the total egg-laying rate is significantly decreased for *D. suzukii* when aged on standard medium (n=50, 30, 30, 50, 50). We thus performed all subsequent experiments with flies raised on Nutrifly medium which elicits an equivalent egg-laying rate in both species, unlike the standard medium. **b** No-choice stimulation assays on either ripe or fermented substrates. The fermented substrate is not repellent to *D. suzukii.* For each species, the ripe and fermented substrates stimulate egg-laying to a similar extent (n=20, 20, 20, 20).

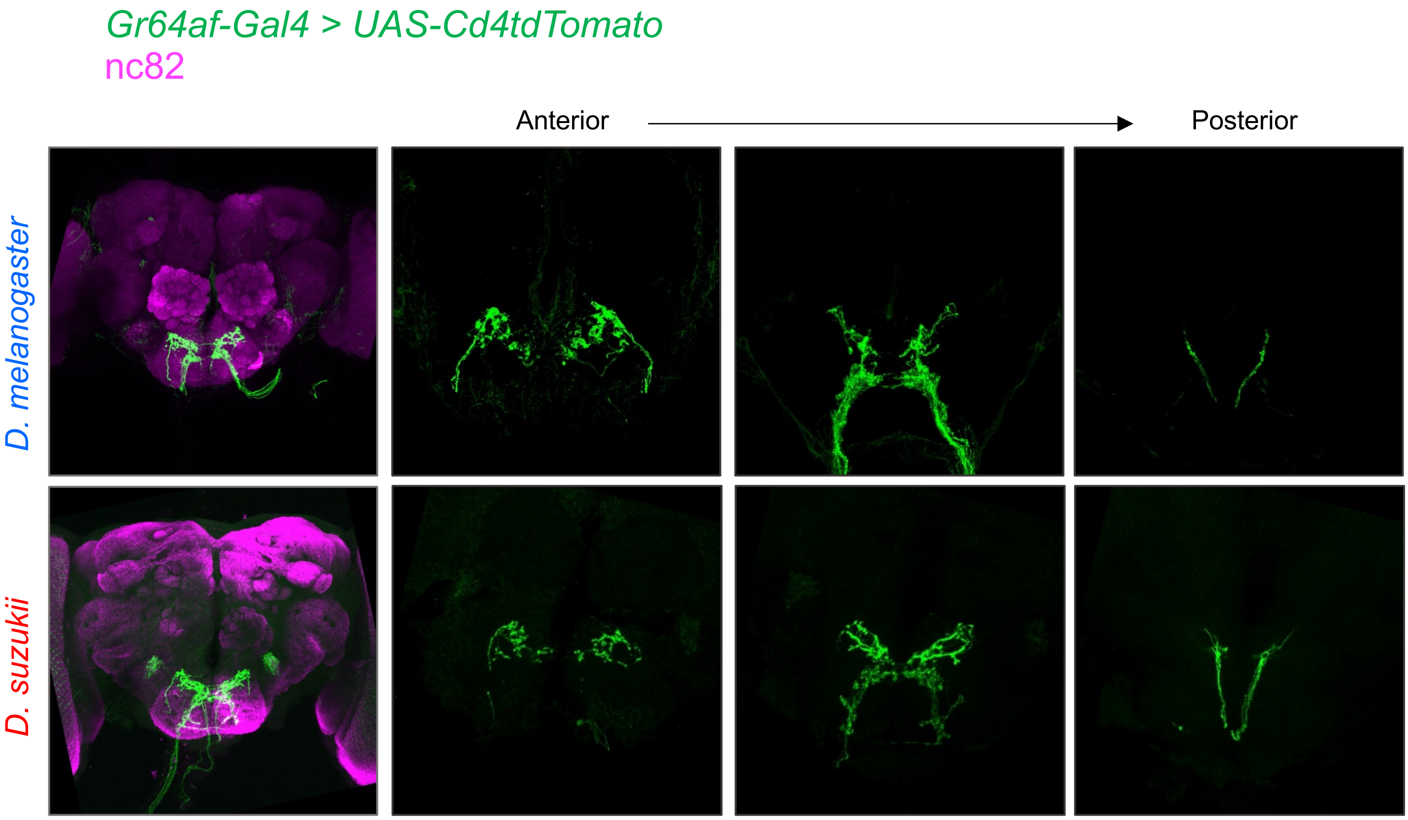


**Supplementary Fig. 2.** *D. suzukii Gr64af-Gal4* neuronal projections onto the CNS.
Axonal projection patterns of the neurons labeled by the *Gr64af-Gal4* lines from *D. melanogaster* (top row) and *D. suzukii* (bottom row) on the CNS. A *UAS-Cd4tdTomato* reporter was used in both species, neuropil stained with nc82 antibody (magenta). *Gr64af-Gal4*-positive neurons project exclusively to the gustatory center – the Sub Esophageal Zone - in both species. Merged images (left-most) show z-projections over all planes. Images on the right and in the green-channel-only show z-projections at different depths from anterior to posterior, revealing similar categories of arborization patterns in both species.

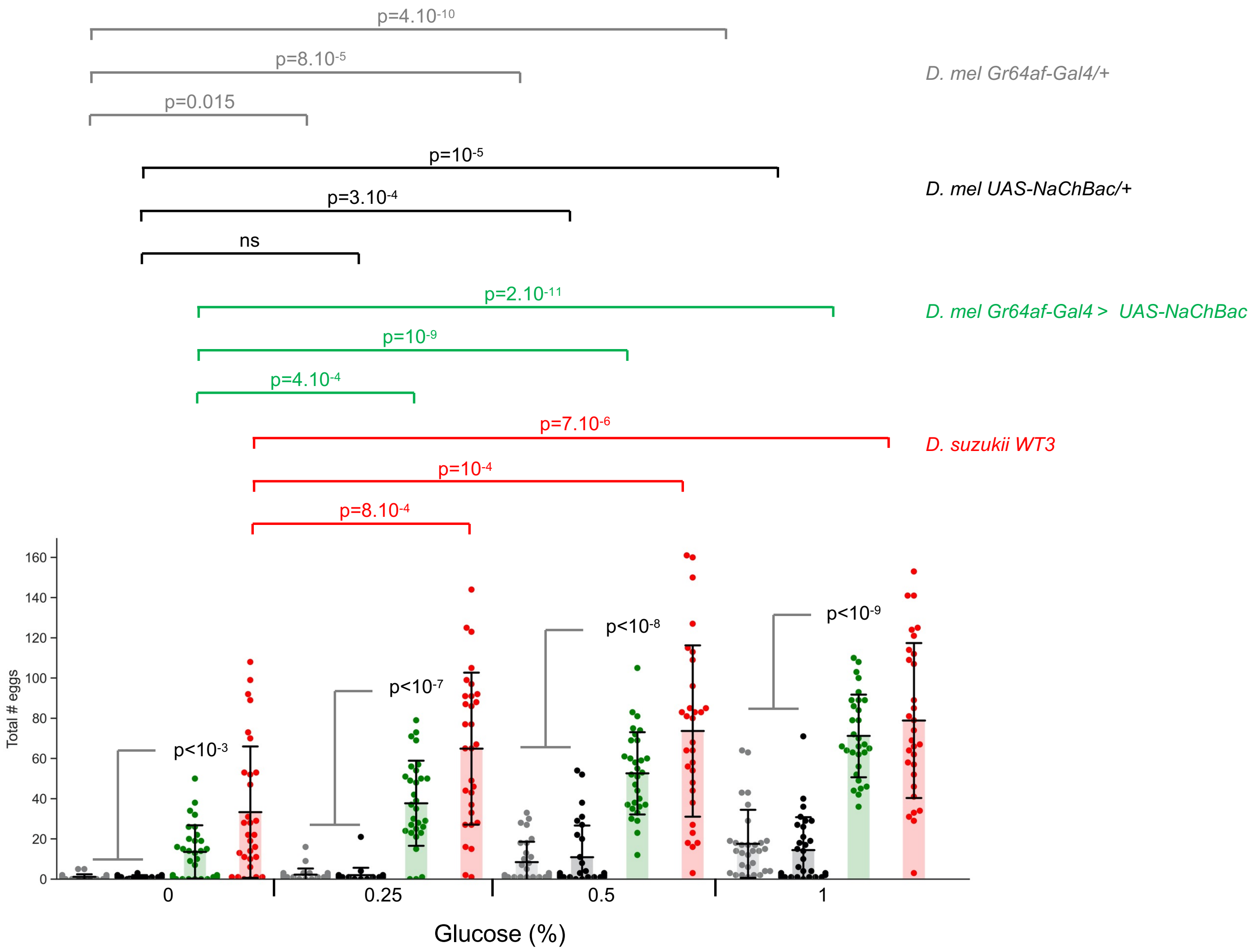


**Supplementary Fig. 3.** Raw data for egg-laying dose-response experiments presented in **Fig. 5a**. Egg-laying rate of the four indicated genotypes in no-choice assays with increasing concentrations of glucose.
